# The stress specific impact of ALKBH1 on tRNA cleavage and tiRNA generation

**DOI:** 10.1101/2020.01.31.928234

**Authors:** Sherif Rashad, Xiaobo Han, Kanako Sato, Eikan Mishima, Takaaki Abe, Teiji Tominaga, Kuniyasu Niizuma

## Abstract

tiRNAs are small non-coding RNAs produced when tRNA is cleaved under stress. tRNA methylation modifications has emerged in recent years as important regulators for tRNA structural stability and sensitivity to cleavage and tiRNA generation during stress, however, the specificity and higher regulation of such a process is not fully understood. Alkbh1 is a m^1^A demethylase that leads to destabilization of tRNA and enhanced tRNA cleavage. We examined the impact of Alkbh1 targeting via gene knockdown or overexpression on B35 rat neuroblastoma cell line fate following stresses and on tRNA cleavage. We show that Alkbh1 impact on cell fate and tRNA cleavage is a stress specific process that is impacted by the demethylating capacity of the cellular stress in question. We also show that not all tRNAs are cleaved equally following Alkbh1 manipulation and stress, and that Alkbh1 KD fails to rescue tRNAs from cleavage following demethylating stresses. These findings shed a light on the specificity and higher regulation of tRNA cleavage and should act as a guide for future work exploring the utility of Alkbh1 as a therapeutic target for cancers or ischemic insult.

## Introduction

tiRNAs are novel small non-coding RNAs produced via tRNA cleavage during stress^1^. tiRNAs were shown to be related to several diseases and pathologies as well as several cellular stressors^2-4^. tiRNAs were shown to repress protein translation and localize to stress granules^5, 6^. Moreover, Angiogenin mediated tRNA cleavage was shown to regulate stem cell function^7^. In recent years, tRNA was revealed to be a rich hub of modifications, especially methylation modifications^1, 8^. Several tRNA methyltransferases were shown to regulate tRNA stability and susceptibility to cleavage under stress such as Nsun2, Dnmt2^9, 10^ and Gtpbp3^11^.

1-methyladenosine (m^1^A) modifications in tRNA occur at positions 9, 14, 22, 57 and 58 in cytosolic tRNA species and at 9 and 58 positions in mitochondrial tRNA species^12^. While they are common in tRNA, m^1^A modifications are comparatively rare in mRNA^13, 14^. m^1^A modifications were shown to impact structural integrity of tRNA, especially those at position 9 and 58, which occurs in both cytosolic and mitochondrial tRNA^15^. Alkbh1 (alkB homolog 1, histone H2A dioxygenase) is a DNA repair enzyme that was shown to regulate several modifications in cytoplasmic and mitochondrial tRNAs^16, 17^. In particular, it was shown to demethylate m^1^A at position 58 in Hela cells^17^ and to be involved in the biogenesis of m^5^C modifications in cytosolic tRNA-Leu and mitochondrial tRNA-Met in HEK293 cells^16^. Interestingly, Alkbh1 overexpression led to translational repression of protein synthesis^17^, while its deletion impaired mitochondrial function^16^. In addition to these somewhat conflicting results on the role of Alkbh1 in cell growth and energy regulation, Alkbh1 did not exhibit its demethylase activity on m^1^A in patient derived glioma stem cells (GSCs)^18^. Notwithstanding, its knockdown in GSCs inhibited cell growth via its DNA N^6^-methyladenine demethylation activity^18^, in contrast to the lower levels of Alkbh1 observed in larger and more advanced gastric cancers indicating an opposing functional roles in both cancer types^19^. Alkbh1 was further shown to localize to mitochondrial RNA granules, exerting an important regulatory function on mitochondrial dynamics^20^. Its knockdown retarded cell growth of Hela cells and delayed development in C. elegans^20^. Alkbh1 was also shown to be of paramount importance for neural and embryonic stem cells differentiation and Alkbh1 homozygous knockout mice die during their embryonic stages^21, 22^. These findings imply an important epigenetic regulatory function of Alkbh1, that appears to be somewhat cell-type related. Since Alkbh1-mediated tRNA demethylation is proposed to impact tRNA stability and increase cleavage, we wanted to evaluate whether modulation of Alkbh1 can impact cytosolic and/or mitochondrial tRNA cleavage, whether knockdown or overexpressing Alkbh1 in neuronal cells can affect cellular fate following different stresses, and whether this can be exploited as a therapeutic strategy for managing oxidative stress induced neuronal apoptosis.

## Results

### tRNA is cleaved following different cell stresses

First, we evaluated tRNA cleavage in B35 neuroblastoma cells following different oxidative stresses. We exposed the cells to increasing concentrations of sodium meta-Arsenite (thereafter termed As), which induces both apoptosis and necrosis via different mechanisms for 6 hours^23^, to Antimycin A, which induces cell death via inhibiting mitochondrial complex III activity leading to reactive oxygen species (ROS) accumulation for 6 hours^24^, and we exposed the cells to 16 hours of oxygen glucose deprivation (OGD) with or without 1 hour of reperfusion as per our protocol^2^. Indeed, As led to robust tRNA cleavage with increasing concentrations as we reported previously with PC12 cells^25^. Antimycin also led to robust tRNA cleavage and tiRNA generation. OGD alone did not induce significant tRNA cleavage, however, 1 hour of reperfusion following OGD (OGD-R) led to significant tRNA cleavage (Figure 1A). Interestingly, when we performed Immuno-northern blotting (INB) with anti-m^1^A antibody we detected only m^1^A harboring tiRNA fragments following As stress. Antimycin stress, on the other hand, showed only minimal tiRNAs harboring m^1^A as compared to As stress, despite a very robust cleavage observed with SYBR gold staining. Interestingly, antimycin caused the decrease in m^1^A signal in the full length tRNA, indicating a demethylation event not observed with As. OGD-R as well showed no m^1^A harboring tiRNA fragments, while leading at the same time to reduction in the m^1^A signal in the full length tRNA itself (Figure 1B).

**Figure 1:**
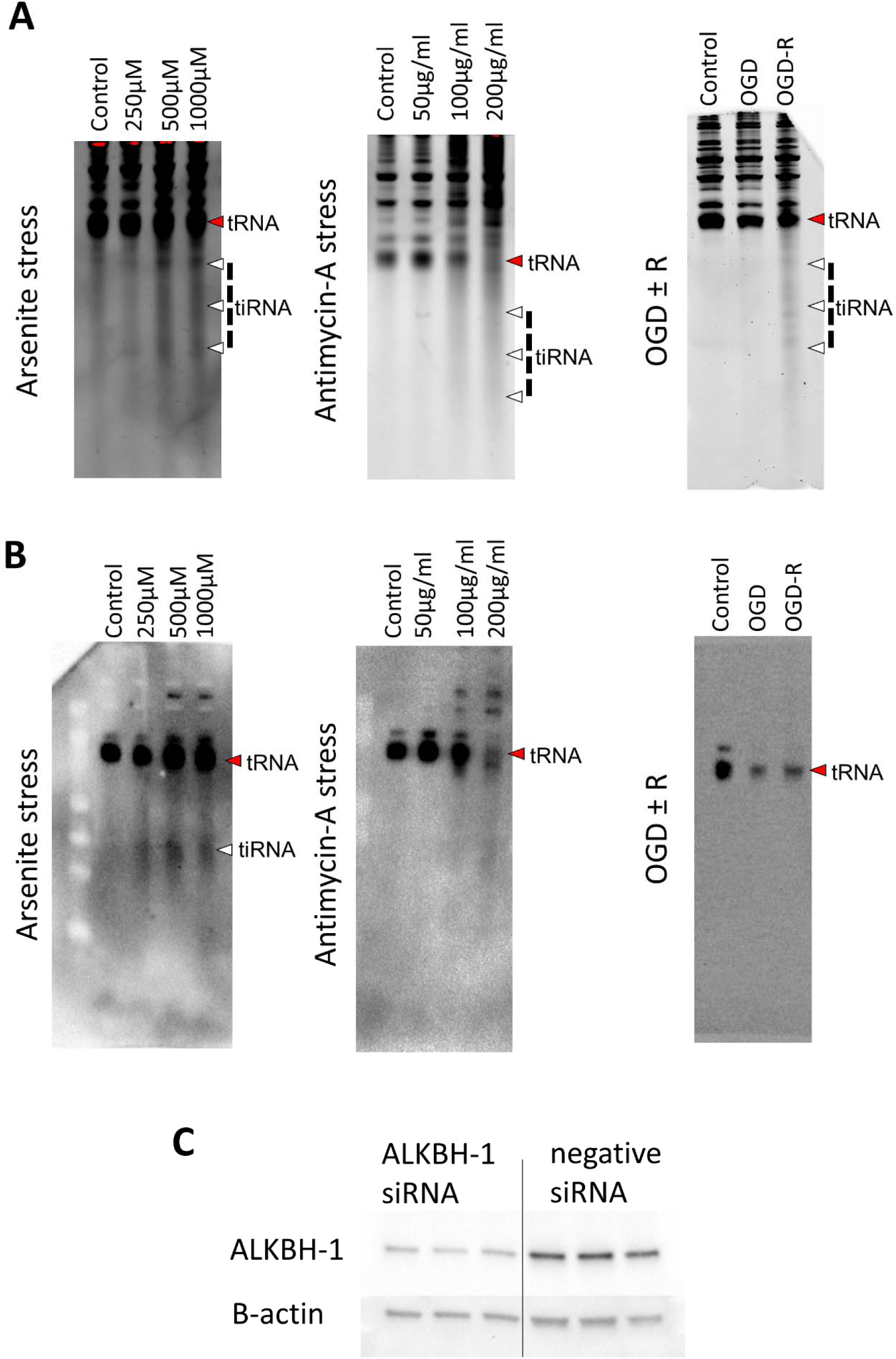
tRNA is cleaved after stress. **A:** B35 cells were exposed to increasing concentrations of arsenite (As), antimycin A for 6 hours or to 16 hours of oxygen-glucose deprivation (OGD) and OGD with 1 hour of reperfusion (OGD+R). SYBR gold staining was performed following electrophoresis of 2µg RNA per well and showed a correlation between the stress level and tRNA cleavage in all stresses [Cropped membranes are shown as a whole in supplementary figure 1]. **B:** Immunonorthern blotting (INB) using anti-m^1^A antibody showed that while following arsenite stress 3’tiRNA fragments harboring m^1^A were detected, antimycin stress showed much lower levels of these fragments. OGD-R showed no m^1^A containing 3’tiRNA fragments. Both antimycin and OGD-R caused reduction of m^1^A levels in full length tRNA as well, something that was not observed to that extent following arsenite stress [Cropped membranes are shown as a whole in supplementary figure 2]. **C:** Western blotting analysis to confirm the effectiveness of Alkbh1 knockdown following siRNA. Alkbh1 protein levels were reduced to about 40-50% of the baseline levels as compared to scramble (negative) siRNA.

### Knockdown of Alkbh1 leads to stress-specific effects

We next examined the impact of Alkbh1 siRNA-based transient knockdown (KD) (Figure 1C) on cell fate and apoptosis using Annexin V flow cytometry assay and on tRNA cleavage. First, Alkbh1 KD significantly improved both apoptosis and necrosis after As stress with concomitant reduction of tRNA cleavage to near control levels as observed with SYBR gold staining (Figure 2A). Alkbh1 KD did not however improve cell fate following antimycin stress and tRNA cleavage levels were nearly the same when compared with negative (scramble) siRNA (Figure 2B). Alkbh1 KD had no impact on cell death following ODG-R, while tRNA cleavage levels were modestly increased as compared to negative siRNA (Figure 2C).

**Figure 2:**
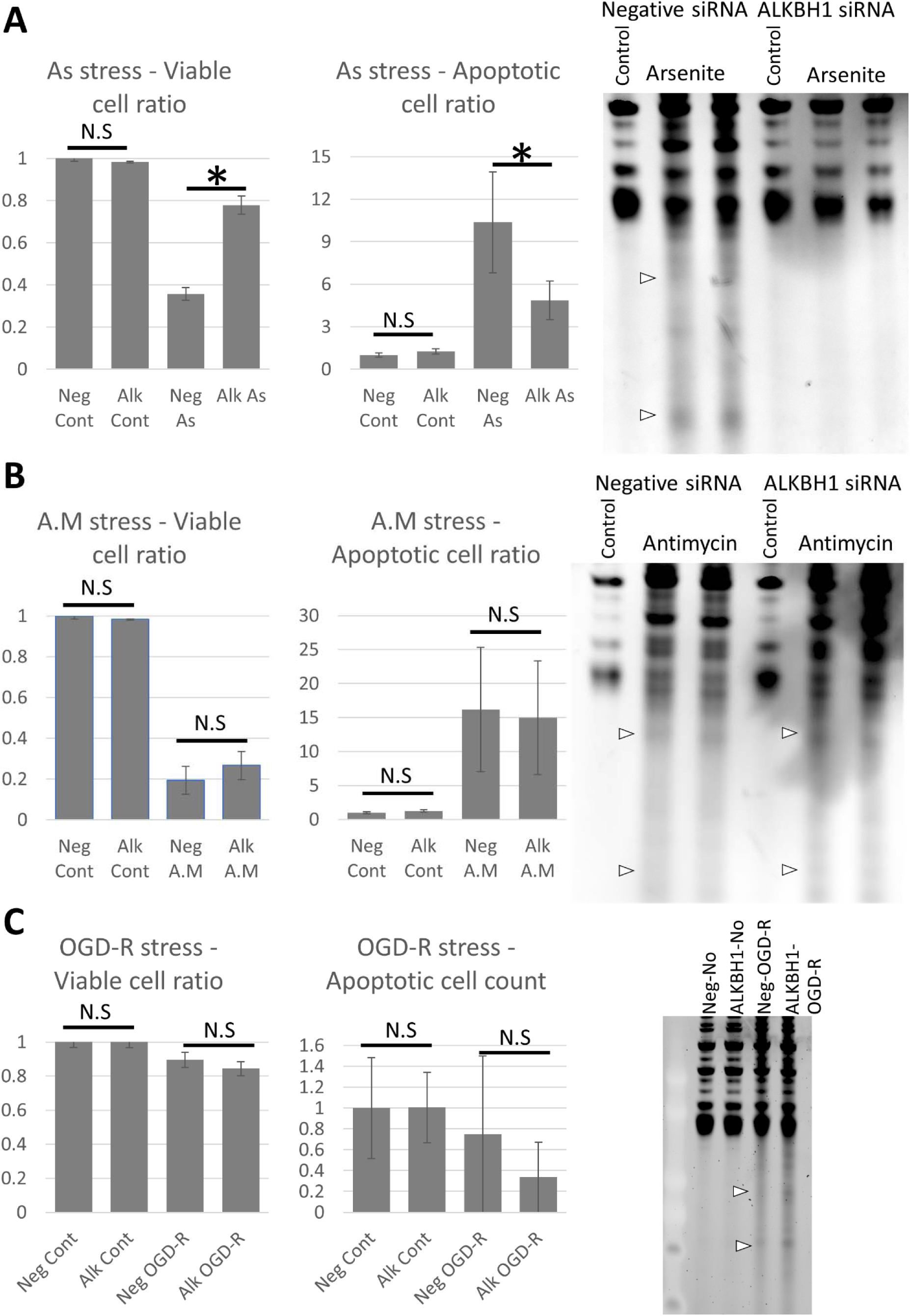
Impact of Alkbh1 knockdown on cell fate and tRNA cleavage after stress. **A:** Annexin V FACS analysis (left graph) and SYBR gold staining following Alkbh1 KD and arsenite stress (400µM for 4 hours). Alkbh1 KD reduced the levels of apoptotic and necrotic cells and rescued tRNA from cleavage. **B:** Annexin V FACS analysis and SYBR gold staining following Alkbh1 KD and antimycin stress (150µg/ml for 4 hours). Alkbh1 KD had no impact on cell fate or tRNA cleavage following antimycin stress. (Note: FACS for As and antimycin stresses were performed simultaneously but the results presented in 2 graphs for easier comprehension and data presentation. FACS data were presented as ratio to unstressed (control) cells). **C:** Annexin V FACS analysis and SYBR gold staining following Alkbh1 KD and OGD-R stress. Alkbh1 KD also had no impact on cell death ratio following OGD-R and no apparent impact on tRNA cleavage. Asterisk: *p* < 0.05, N.S: not significant statistically. SYBR gold staining was performed with 500ng total RNA per lane.

### Impact of Alkbh1 KD on tRNA cleavage following different stresses

To further evaluate tRNA cleavage, we probed different tRNAs, cytosolic (Figure 3A) and mitochondrial (Figure 3B), using DIG-labeled probes against the 5’tiRNA halves, focusing on As and antimycin stresses, since the stress duration is similar which allows for easy comparison between both stresses, after Alkbh1 siRNA-based transient knockdown. Certain tRNAs were shown to be bound to Alkbh1 using CLIP^17^, thus, we probed some of these tRNA species as well as other non-Alkbh1 bound tRNAs according to Liu et al^17^.

**Figure 3:**
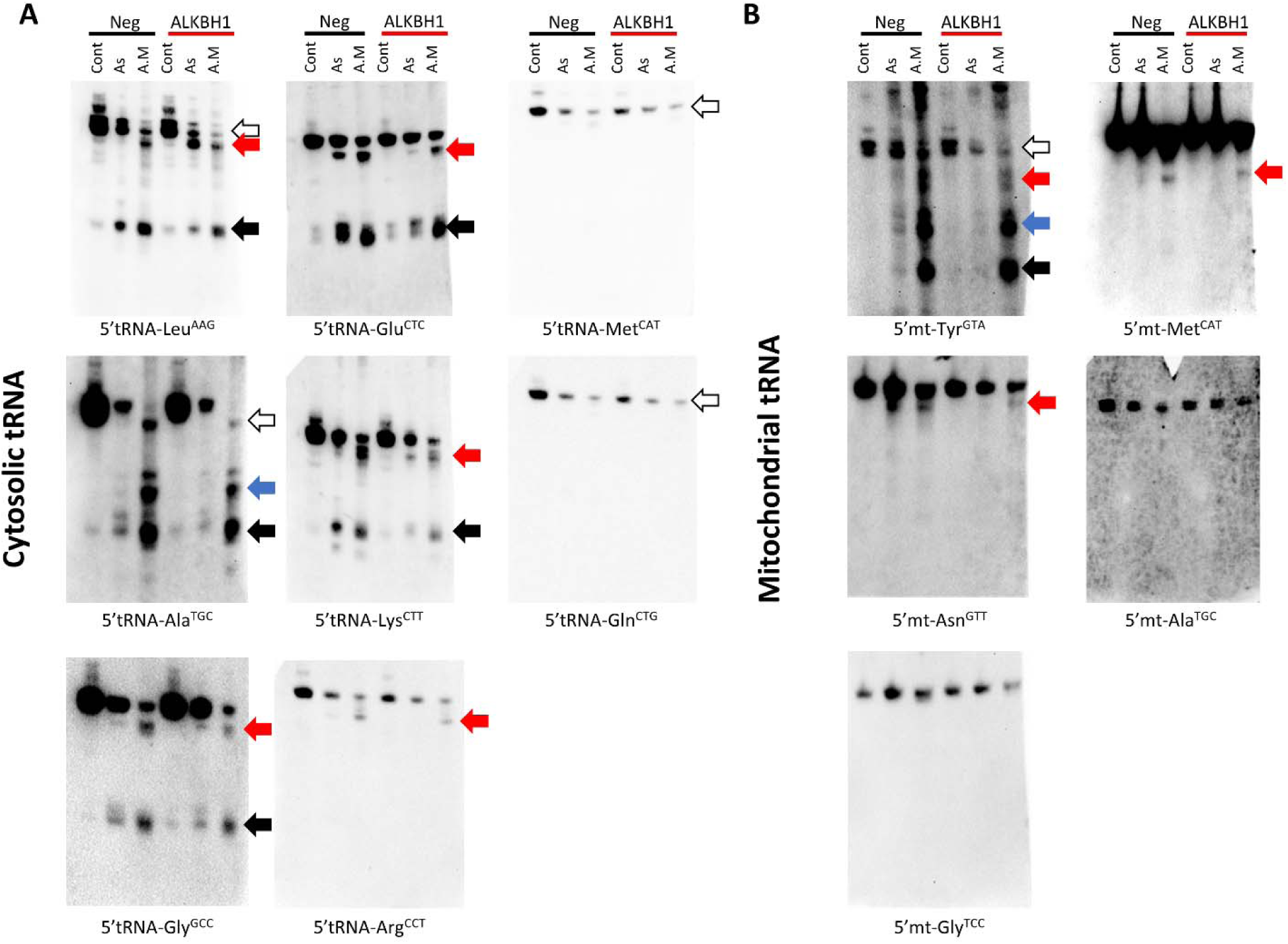
Impact of Alkbh1 KD on different cytosolic and mitochondrial tRNA species’ cleavage show stress and tRNA specific patterns. **A:** Cytosolic tRNA cleavage before and after Alkbh1 KD following 4 hours of arsenite (As; 400µM) or antimycin (A.M; 150µg/ml) stresses. Northern blotting performed with 1µg total RNA per lane. Leu^AAG^ 5’tiRNA cleavage was significantly reduced after Alkbh1 KD following As stress, with little to no effect on its cleavage following A.M stress. There was a notable but unexplained reduction in the full length tRNA levels following KD and A.M stress. Moreover, an extra band appeared after As cleavage and KD just below the full length tRNA (similar to one observed with A.M stress) which cannot be explained at the present time (black arrow: 5’tiRNA, white arrow: full length tRNA, Red arrow: large 5’tiRNA fragments). Ala^TGC^ was cleaved into several 5’tiRNAs of varying lengths especially after A.M stress (Blue and black arrows). KD of Alkbh1 reduced the cleavage after As but not A.M stress. Again, the full length tRNA levels were reduced with KD following A.M stress and there is an apparent shift in the tRNA size between A.M and As and control samples. This may indicate partial cleavage of tRNA (white arrow). Gly^GCC^ was cleaved into 2 distinct 5’ fragments, one at the usual size of the 5’tiRNA (black arrow), and another large fragment around 50∼60 nt in length (red arrow). Alkbh1 KD reduced Gly^GCC^ cleavage after As stress, while after A.M stress the larger fragments showed a decrease in signal while the canonical fragments (black arrow) were not significantly changed. Glu^CTC^ also showed cleavage into 2 5’tiRNA fragments. Alkbh1 KD impacted the cleavage after As stress only. Lys^CTT^ was also cleaved into 2 5’tiRNA fragments. Alkbh1 KD reduced the cleavage following As and A.M stresses. Arg^CCT^ cleavage was apparent in the form of large 5’tiRNA fragment after A.M stress only, and this fragment was minimally affected by Alkbh1 KD (although this effect can be negligible). Met^CAT^ and Gln^CTG^ did not exhibit 5’tiRNA fragments after stress, but there was a reduction in their full length tRNA levels (white arrows). Alkbh1 had no apparent impact on these 2 tRNAs. **B:** Mitochondrial tRNA cleavage before and after Alkbh1 KD following 4 hours of arsenite (As; 400µM) or antimycin (A.M; 150µg/ml) stresses. Northern blotting performed with 4µg total RNA per lane. Mt-Tyr^GTA^ was cleaved into several fragments after As and A.M cleavage, with more robust cleavage after A.M stress. Alkbh1 KD reduced mt-Tyr^GTA^ cleavage after As but not A.M stress. Notably, the largest of the 5’tiRNA fragments (red arrow) was reduced with Alkbh1 KD following A.M stress. Moreover, full length tRNA levels (white arrow) were reduced after KD. mt-Asn^GTT^ was cleaved into a larger 5’tiRNA fragment (red arrow) which was reduced with Alkbh1 KD following As and A.M stresses. mt-Met^CAT^ was cleaved into a large 5’tiRNA fragment as well apparent only after A.M stress. Alkbh1 KD minimally reduced the signal of the 5’tiRNA-mt-Met^CAT^. mt-Gly^TCC^ and mt-Ala^TGC^ were not apparently cleaved and showed no impact of Alkbh1 on their levels. [Membranes were not cropped]

First, tRNA-Leu^AAG^, a major substrate for Alkbh1^16, 17^ was robustly cleaved with both As and antimycin stresses, however, Alkbh1 knockdown reduced the degree of cleavage after As stress, while it had no major impact on the antimycin induced cleavage.

Ala^TGC^, another Alkbh1 interacting tRNA^17^, showed the same response as with Leu^AAG^. Interestingly, Ala^TGC^ was cleaved into more than one fragment with distinct nucleotide lengths. Moreover, Alkbh1 siRNA not only failed to significantly reduce the cleavage of tRNA-Ala^TGC^ following antimycin stress, it reduced the levels of the near-full length Ala^TGC^ as well as the larger 5’tiRNA fragments.

Gly^GCC^, also a substrate of Alkbh1, was also cleaved with both stresses and the cleavage also was into 2 distinct fragments, with the larger fragment more evidently observed with antimycin stress. Alkbh1 knockdown did reduce the signal of the larger fragment with antimycin stress, however the smaller fragment showed no significant change. There was also a modest reduction of the 5’tiRNA signal after As stress, but not as evident as with the previous 2 species.

Glu^CTC^, another Alkbh1 substrate, was also cleaved into 2 distinct 5’tiRNA fragments. Alkbh1 KD reduced its cleavage after As stress but not after antimycin stress. Lys^CTT^, a major Alkbh1 substrate^17^, also showed the same cleavage pattern into 2 distinct 5’tiRNA fragments. Alkbh1 KD reduced the cleavage into either fragment following As and antimycin stresses.

tRNA-Arg^CCT^, which is not a physically interacting tRNA with Alkbh1^17^, was cleaved into a large 5’tiRNA fragment after antimycin stress only. Alkbh1 KD did not significantly impact the level of this fragment.

Gln^CTG^, another tRNA that binds to Alkbh1, did not show a 5’tiRNA fragment after As or antimycin stress, rather the full length tRNA itself was reduced. Contrary to other Alkbh1 bound tRNAs, Alkbh1 KD did not affect Gln^CTG^ significantly after stress.

tRNA-Met^CAT^ showed a similar pattern as with Gln^CTG^ regarding the reduction of full length tRNA levels. Alkbh1 knockdown was shown previously to increase the levels of Met^CAT 17^, However, we did not observe any significant changes in Met^CAT^ following Alkbh1 KD before or after stress. Overall, we did not observe significant 5’tiRNA fragments for Met^CAT^ or Gln^CTG^. This might be due to a very low levels of these fragments, below the threshold of detection by northern blotting. Nonetheless, we cannot make assumptions on the changes in the cleavage of these tRNAs after stress aside from the inference we got from observing the full length tRNA levels.

As for mitochondrial tRNAs (Figure 3B), Alkbh1 KD reduced the global cleavage of the mt-tRNA-Tyr^GTA^ after As stress. After antimycin toxicity, mt-Tyr^GTA^ was cleaved into a number of distinct 5’tiRNA fragments of various lengths (observed also with As but to a lesser extent). Alkbh1 KD reduced the levels of the full length mt-tRNA-Tyr^GTA^ itself as well as the largest of the cleaved fragments, while the smaller 5’tiRNA fragments did not show changes in their levels.

mt-Asn^GTT^ was cleaved into a large 5’tiRNA fragment that is very close in size to the full length tRNA after As and antimycin stresses. Alkbh1 KD protected this tRNA from cleavage after As stress and to a lesser extent after antimycin stress.

The mitochondrial tRNA-Met^CAT^, contrary to its cytosolic counterpart, showed a cleaved 5’tiRNA fragment after antimycin stress, and not As stress. Again, this fragment was of a larger size than the canonical 35∼45nt 5’tiRNA fragments. Alkbh1 KD did not significantly affect the levels of its cleavage and only minimal reduction of its signal was observed.

mt-Gly^TCC^ and mt-Ala^TGC^ did not exhibit significant changes or cleavage after stress with or without Alkbh1 KD.

### Impact of Alkbh1 over expression on cell fate and tRNA cleavage

To further confirm our data, we stably overexpressed Alkbh1 in B35 cells using Lenti-viral based transduction and single colony selection and expansion, which resulted in robust expression of exogenous rat Alkbh1 in the cells in the total cellular protein extract (Figure 4A). Control lentiviral particles encoding a scramble sequence were also transduced into B35 cells followed by single colony expansion. Alkbh1 over expression cells are referred to as B35^ALK^, while control cells are referred to as B35^Mock^. Both cell lines showed comparable growth as evident by WST-8 assay and repeated cell counting in 6 well plate (Supplementary figure 3). Alkbh1, similar to our observations with the KD, affected cell survival following As stress and not antimycin stress, leading to more cell death and increased tRNA cleavage following As stress (Figure 4B & C).

**Figure 4:**
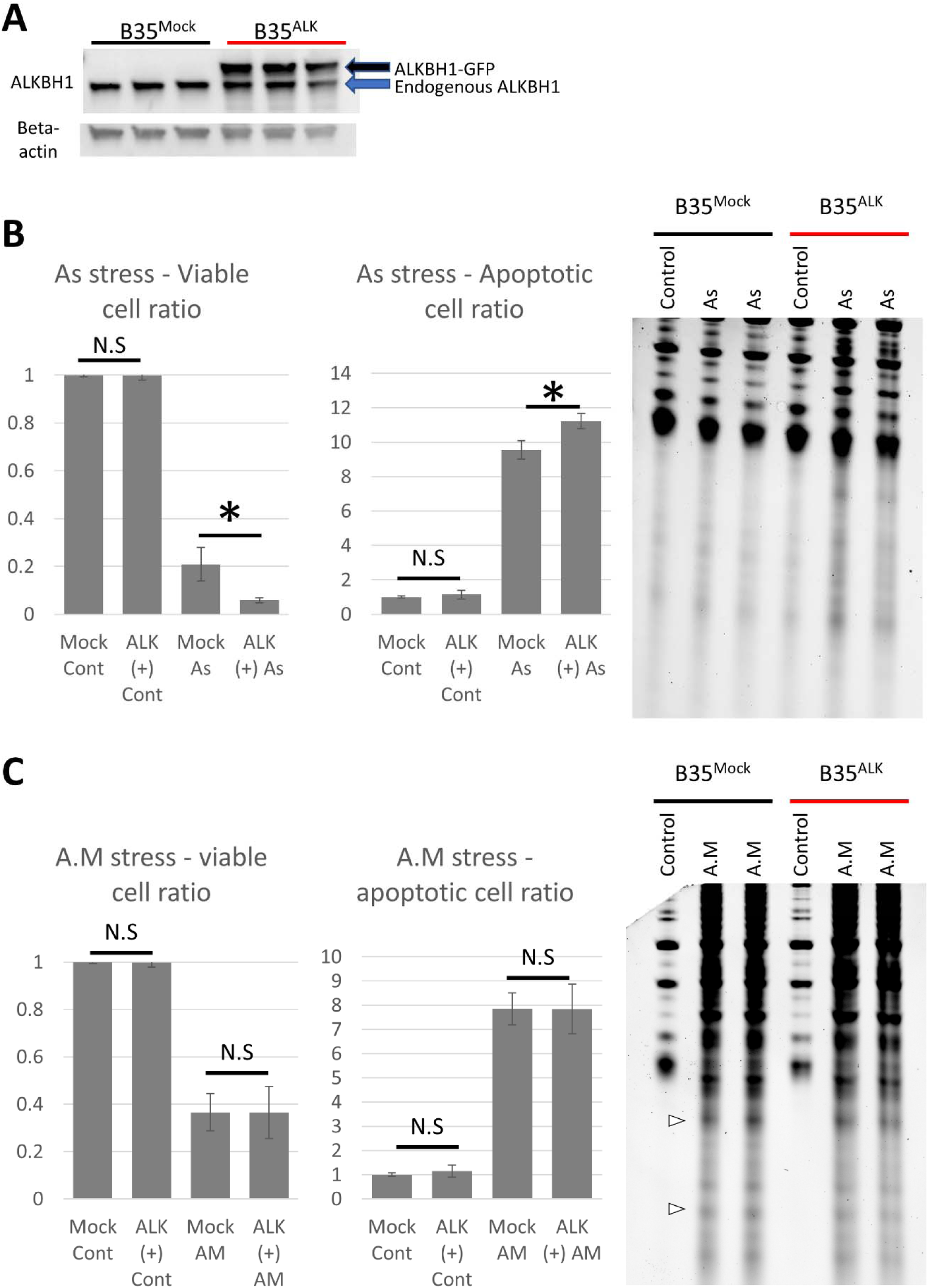
Impact of Alkbh1 overexpression on cell fate and tRNA cleavage. **A:** Western blotting confirmation of Alkbh1 overexpression. Lenti-encoded Alkbh1 with GFP-attached sequence resulted in a larger protein (black arrow) than the endogenous Alkbh1 (blue arrow). **B:** Arsenite stress (4 hours, 1mM) resulted in more cell death in B35^ALK^ than in B35^Mock^ cells as revealed by Annexin V FACS assay (left graphs). This effect was reflected in the form of increased gross tRNA cleavage and tiRNA generation observed on SYBR gold staining (Right). **C:** Antimycin stress (4 hours, 200µg/ml) resulted in equal levels of cell death in B35^Mock^ and B35^ALK^ (left: Annexin V FACS analysis) and equal levels of tRNA cleavage (Right: SYBR gold staining). (Note: FACS for As and antimycin stresses were performed simultaneously but the results presented in 2 graphs for easier comprehension and data presentation). Asterisk: *p* < 0.05, N.S: not significant statistically. SYBR gold staining was performed with 500ng total RNA per lane. FACS data were presented as fold-change unstressed cells.

Alkbh1 overexpression also enhanced the cleavage of several tRNA species in a stress specific behavior, in a manner opposing to and confirming what we observed with Alkbh1 KD as shown in Figure 5. Interestingly, Alkbh1 overexpression enhanced the cleavage of Met^CAT^, mt-Met^CAT^, Gln^CTG^ and mt-Asn^GTT^, leading to the observation of previously unobservable 5’tiRNA fragments (Figure 5A & B). Collectively, these results indicate that Alkbh1 impact on tRNA is tRNA species specific as well as in a stress specific. However, Alkbh1 does not seem to impact tRNA cleavage or total tRNA levels in the absence of oxidative stress.

**Figure 5:**
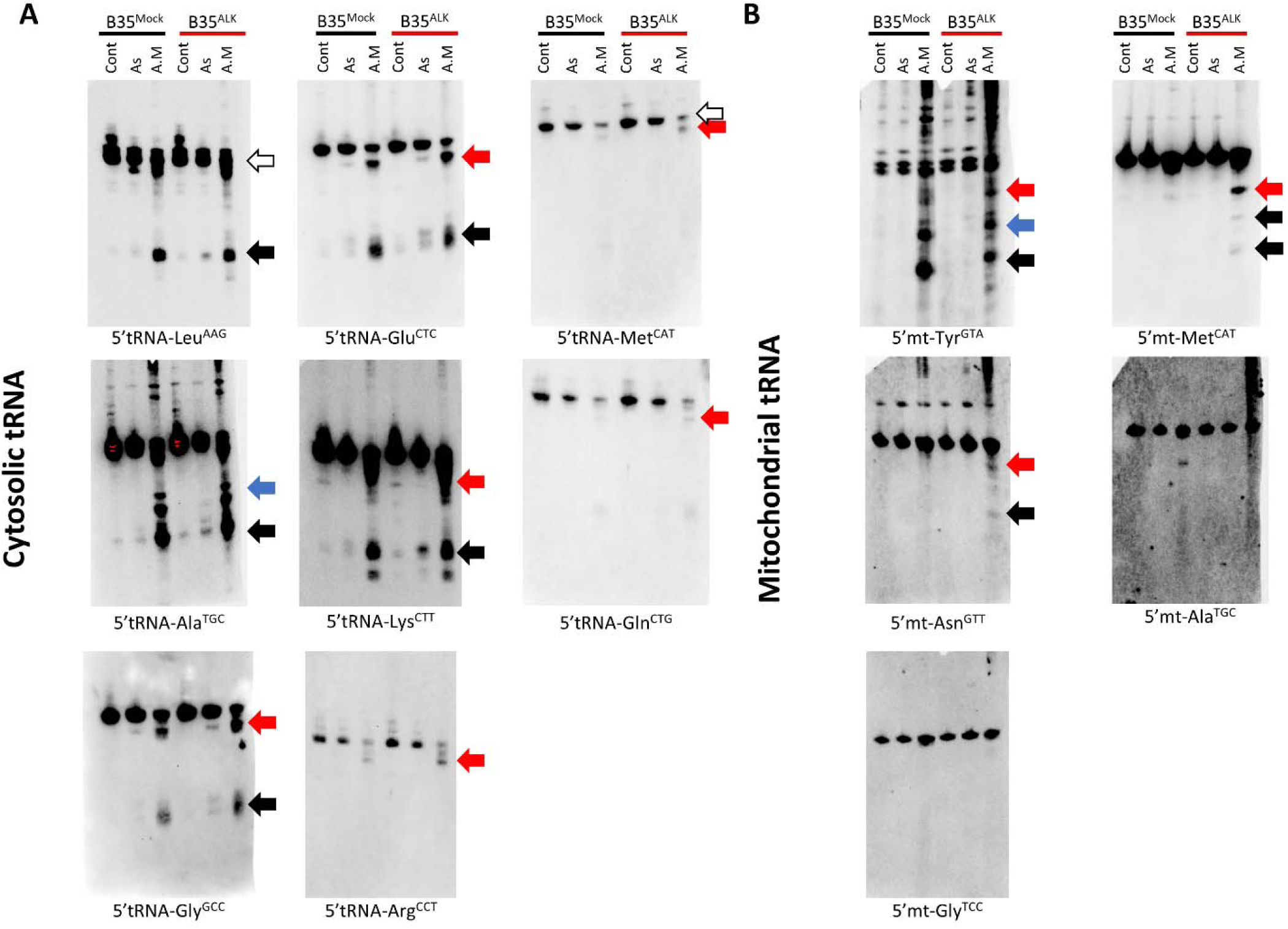
Impact of Alkbh1 overexpression on different cytosolic and mitochondrial tRNA species’ cleavage show stress and tRNA specific patterns. **A:** Cytosolic tRNA cleavage in B35^Mock^ and B35^ALK^ cells after 4 hours of As (1mM) or A.M (200µg/ml) stresses. Northern blotting performed with 1µg total RNA per lane, Alkbh1 overexpression led to a general pattern of increased tRNA cleavage. Leu^AAG^, Ala^TGC^, Gly^GCC^, Glu^CTC^, Lys^CTT^ and Arg^CCT^ showed increased cleavage in their 5’tiRNA fragments (whether canonical or larger fragments) in a pattern opposite to what was observed with Alkbh1 KD (figure 3). Moreover, Met^CAT^ and Gln^CTG^, which did not exhibit apparent cleavage with stress and siRNA KD, were cleaved with Alkbh1 overexpression into large 5’tiRNA fragments after A.M stress (red arrows). **B:** Mitochondrial tRNA cleavage in B35^Mock^ and B35^ALK^ cells after As and A.M stresses. Northern blotting performed with 4µg total RNA per lane. mt-Tyr^GTA^ cleavage after As in B35^ALK^ cells was not impacted in an apparent manner, however, after A.M stress, the larger fragments (red arrow), which were reduced after siRNA KD in figure 3, did exhibit some increase in its signal. mt-Met^CAT^ cleavage after A.M stress in B35^ALK^ cells was greatly enhanced, with the larger fragment (red arrow) being clearly visible and the appearance of 2 other fragments that were not observable prior to Alkbh1 overexpression (black arrows). mt-Asn^GTT^ showed an enhanced cleavage after Alkbh1 overexpression (red arrow), with the observation of faint signal of a smaller 5’tiRNA fragment not observable prior to overexpression (black arrow). mt-Gly^TCC^, mt-Arg^TCG^ and mt-Ala^TGC^ that were not cleaved prior to Alkbh1 overexpression showed no impact of Alkbh1 on their levels.

### Impact of ALKBH1 on tRNA m^1^A methylation after stress

Since Alkbh1 was shown to demethylate m^1^A^17^, we sought to evaluate the impact of Alkbh1 on the tRNA methylation status following KD or overexpression and after stress using INB. Alkbh1 KD resulted in the partial recovery of full length tRNA m^1^A methylation as well as tiRNA reduction following As stress (Figure 6A). Surprisingly, it led to a reduction of tRNA m^1^A status following antimycin stress, contrary to what we expected (Figure 6A). Antimycin appears to be an m^1^A demethylating stressor (Figure 1B), however, the reason for Alkbh1 KD to be acting in synergy with antimycin to enhance tRNA demethylation is not clear.

**Figure 6:**
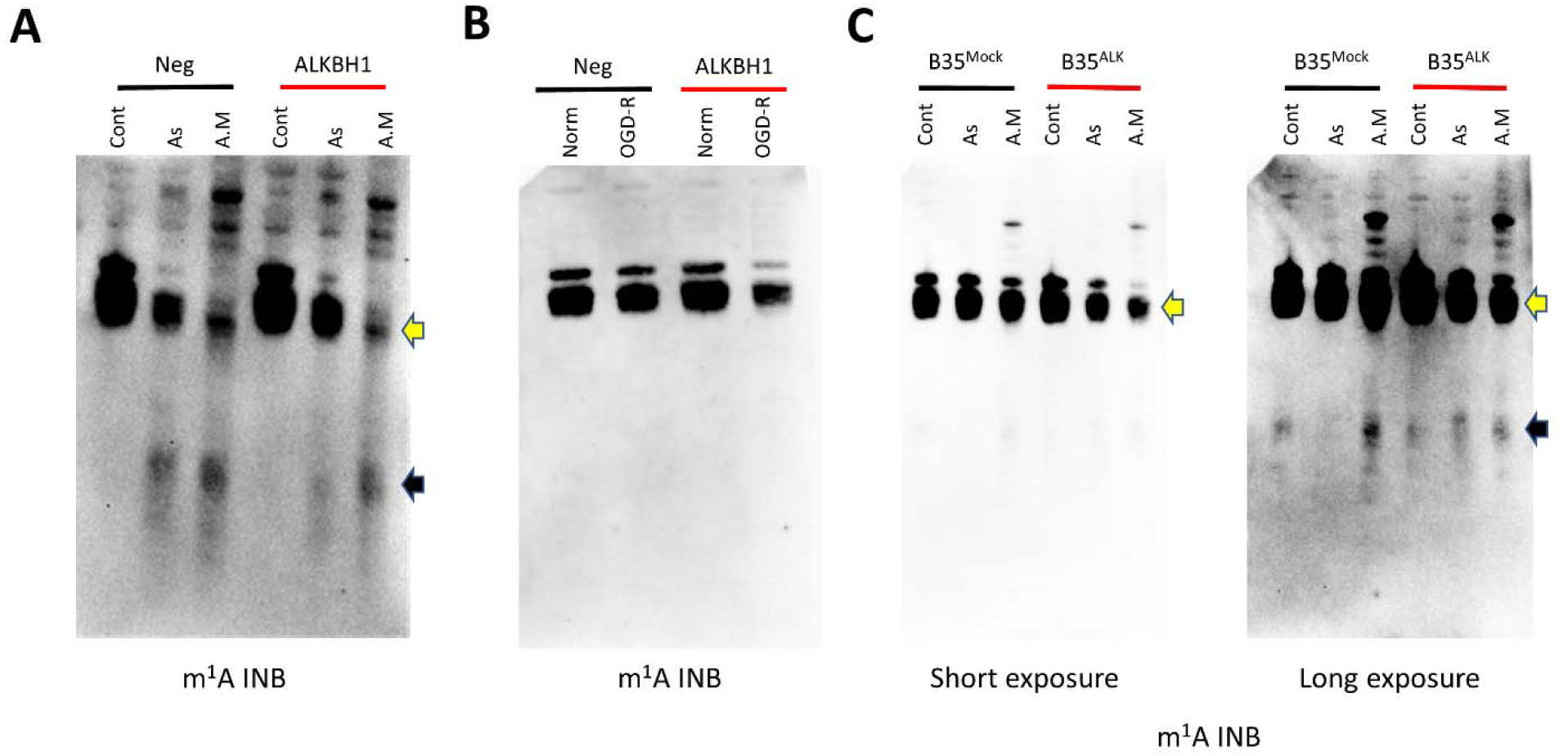
Stress specific impact of Alkbh1 on m^1^A tRNA status. Immunonorthern blotting (INB) with anti-m^1^A antibody following stress and Alkbh1 siRNA KD (**A and B**) or Alkbh1 overexpression (**C**). 4µg total RNA were loaded per lane to enhance the resulting signal. **A:** As and A.M stress both resulted in tRNA cleavage as observed with the existence of m^1^A-harboring 3’tiRNA fragments (black arrow). As and A.M stresses both led to a demethylation effect on tRNA (yellow arrow), however, As stress resulted in a moderate demethylation as opposed to the significant demethylation observed with A.M. Alkbh1 KD resulted in decrease tRNA cleavage and recovery of tRNA m^1^A methylation status after As stress, while after A.M stress there was no impact of Alkbh1 KD on tRNA cleavage and tRNA was further demethylated in a paradoxical manner. **B:** Alkbh1 KD following OGD-R stress resulted in decreases m^1^A signal in full length tRNA indicating a paradoxical demethylating effect. **C:** INB after As (1mM) and A.M (200µg/ml) stress exposure of B35^Mock^ and B35^ALK^ for 4 hours. Alkbh1 overexpression resulted in modest demethylating effect on tRNA after As stress and a significant demethylation after A.M (yellow arrow). Alkbh1 overexpression also led to increase in m^1^A-harboring tiRNAs after As stress and a paradoxical reduction of such fragments after A.M stress indicating that the demethylation also impacts the resulting tiRNAs.

Following OGD-R stress, no m^1^A harboring tiRNAs were observed with or without Alkbh1 KD, despite a robust tRNA cleavage detection with SYBR gold staining (Figure 2C), indicating a strong demethylation event. The tRNA itself showed a decrease in the methylation status following Alkbh1 KD and OGD-R, similar to what was observed with antimycin stress (Figure 6B).

Finally, we examined tRNA m^1^A status in B35 cells with Alkbh1 overexpression after As and antimycin stresses (Figure 6C). Indeed, overexpressing Alkbh1 resulted in tRNA demethylation especially after antimycin stress, albeit modest after As stress. Moreover, Alkbh1 overexpression resulted in more m^1^A-harboring 3’tiRNA fragments to be observed following As stress, however, it led to a paradoxical reduction in these fragments after antimycin stress. This may indicate that the m^1^A demethylation is also reflected on tiRNA fragments and that even though the gross cleavage of tRNA is not much different between B35^Mock^ and B35^ALK^ following antimycin stress, the level of methylated tiRNAs in B35^ALK^ is less. How this demethylation event impacts the function of tiRNA is yet to be determined.

## Discussion

In this work we present several interesting finds. First, we show that tRNA cleavage is not a haphazard process, and the degree, pattern and regulation of different tRNA species’ cleavage is a well-regulated process involving, at least in part, the methylation status of tRNA. We also show that tRNA cleavage patterns depend on the stresses that the cells are exposed to. For instance, the m^1^A demethylating stress caused by antimycin A toxicity leads to more robust and different pattern of tRNA cleavage that non-demethylating arsenite stress. Interestingly, Alkbh1 impact on tRNA cleavage appears to be affected by the degree of tRNA demethylation following different stresses. When the cells were exposed to the demethylating antimycin toxicity or OGD-R stresses, Alkbh1 siRNA mediated knockdown did not impact the cell fate. On the other hand, when tRNA cleavage occurred without severe m^1^A demethylation, as observed in arsenite stress in the form of less demethylation of full-length tRNA and presence of more evident 3’tiRNAs harboring m^1^A, Alkbh1 KD did improve cellular responses and reduce apoptosis. Indeed, Alkbh1 is not the sole tRNA m^1^A demethylase. Recently, Alkbh3, another member of the same family, was shown to also demethylate m^1^A and impact tRNA stability^26^. However, CLIP-seq data showed a different affinity patterns for tRNA species for both enzymes^17, 26^. Moreover, other tRNA modifications impact stability of tRNA and were not addressed in our work^11, 27, 28^. Therefore, we expect, at least in part, that the differences in the effect of Alkbh1 KD on different stresses is due to different dynamics of tRNA methylation/demethylation and the enzymes involved in these processes and how they interact with tRNA in different stress conditions. An important implication of these findings is that therapies directed towards Alkbh1, for example as proposed for glioma^18^, should take into consideration the tRNA methylation status changes with different therapies, even if tRNA appears not to be demethylated by Alkbh1 under quiescent conditions^18^.

It is important to stress that we did not observe a significant change in tRNA m^1^A status following Alkbh1 overexpression at rest, despite a significant impact of Alkbh1 overexpression or KD on tRNA methylation and cleavage after stress. Indeed, our methodology to detect m^1^A using immunonorthern blotting^29^ is less sensitive when compared to mass spectrometry based methods^17^. Nevertheless, we also did not observe changes in total tRNA levels of tRNA-Met as observed by Liu et al^17^ in the absence of stress, which might indicate not only a stress specific impact of Alkbh1 on tRNA but also a cell specific impact, something that should be taken in consideration when designing therapies or studying the tRNA demethylating effect of this enzyme. Interestingly, Alkbh1 had a greater impact on mitochondrial Met^CAT^ than on its cytosolic counterpart. This is inline with the fact that Alkbh1 has an important mitochondrial regulatory function^20^ and that its deletion impaired mitochondrial activity^16^. Alkbh1 was also shown to be involved in the m^5^C modification biogenesis in mt-Met^CAT 16, 30^. Combined with the observed impact on mt-Met^CAT^ cleavage after antimycin induced mitochondrial stress, it is clear that Alkbh1 plays an important and central role in regulating mitochondrial dynamics and responses to stress. It is important to note that Kawarada et al^16^ reported a site and tRNA specific m^1^A demethylase activity of Alkbh1 on mitochondrial tRNAs, further confirming the specificity of Alkbh1 action on tRNA.

While our results stress that m^1^A modifications has a great impact on tRNA stability^17^, other modifications such as m^5^C also impact tRNA stability^9, 10, 31^, the redundancy of these modifications in determining tRNA sensitivity to cleavage needs to be studied and well characterized in a cell/stress specific manner as well as in a tRNA species specific manner.

In conclusion; we show that the impact of Alkbh1 on tRNA is a complex process, regulated, at least in part, by cellular responses to different stresses. We can also state that tRNA cleavage is not homogenous amongst different tRNA species, in terms of cleavage site, and it is a stress specific process as well. The utility of Alkbh1 as a therapeutic target should be evaluated on a case-by-case basis, as a universal approach to its function will be simplistic and will fail to appreciate the complex nature of its function. Finally, the redundancy of m^1^A for tRNA stability and cleavage should be evaluated and characterized, and other m^1^A demethylases and well as methyltransferases should be identified and studied in the context of stress induced tRNA cleavage.

## Methods

### Cell line and materials

B35 neuroblastoma cells were purchased from ATCC (Cat# CRL-2754, RRID: CVCL_1951, Lot# 59301600). Rat Alkbh1 siRNA was purchased from Thermo Fischer (Silencer select siRNA, Cat# 4390771, siRNA ID: s169728). Negative control siRNA was purchased from Thermo Fischer (Silencer select siRNA, Cat# 4390843). Alkbh1 Rat Tagged ORF Clone Lentiviral Particles were purchased from Origene (Cat# RR214755L2V). Lipofectamine RNAimax was purchased from Thermo Fischer (Cat# 13778150). Polybrene was purchased from Sigma-Aldrich (Cat# TR-1003-G). Sodium (meta)Arsenite was purchased from Sigma-Aldrich (Cat# S7400-100G, Lot# SLBR8768V). Streptococcal antimycin A was purchased from Sigma-Aldrich (Cat# A8674, Lots # 067M4074V and 039M4107V).

### Antibodies used

anti-1methyladenosine antibody (1:1000 dilution, MBL, Nagoya, Japan, clone AMA-2, Cat # D345-3); anti-Alkbh1 antibody (1:500 dilution, Abcam, Cat# ab195376); anti-beta actin antibody (1:5000 dilution, Cell signaling technologies, Cat# 4970S); HRP-linked anti-mouse antibody (1:2000, Cell signaling, Cat# 7076S); IgG detector solution, HRP-linked (1:2000, Takara, japan, Cat#T7122A-1);

### Cell culture and cell stress

B35 cells were cultured in high glucose DMEM (Nacalai Tesque, Cat# 08457-55) containing 10% fetal bovine serum (FBS, Hyclone, GE-life sciences, USA. Cat# SH30910.03). For arsenite and antimycin stresses, cells were plated at a density of 1 × 10^6^ in 6-well Poly-L-Lysine coated plates (Corning life sciences, NY. Cat# 354515) or a density of 2 × 10^5^ in Poly-L-Lysine coated 24 well plates (Corning life sciences, NY. Cat# 354414) 1 day prior to the experiment. The next day, growth medium (GM) was changed with medium containing the designated concentrations of arsenite or antimycin A for 6 hours (for wild-type cells) or 4 hours (for siRNA transfected or Alkbh1 overexpressing cells) and then the cell samples collected for further processing. For ischemia-reperfusion stress (I/R) cells were plated the same way and then exposed to 16 hours oxygen-glucose deprivation (OGD) in low glucose DMEM medium (Wako, Cat# 042-32255) with no serum followed by 1 hour of reperfusion in full growth medium as we previously described^2^.

### siRNA knockdown of Alkbh1

Cells were plated at a density of 2 × 10^5^ per well in 6-well plates or 4 × 10^4^ in 24 well plates. The next day medium was changed to high glucose DMEM without FBS or antibiotics and the cells were transfected with Alkbh1 siRNA or Negative control siRNA using Lipofectamine RNAimax according to the manufacturer’s protocol. After 48 hours the medium was changed to full GM and the cells allowed to recover for 24 hours before stress experiments or sample collection for RNA or western blotting analysis. Transfection efficiency was > 50% as evaluated using western blotting (Figure 1C). I/R stress was performed as described previously, however, due to Lipofectamine induced toxicity to the cells, the optimal drug induced oxidative stresses were evaluated to be 400µM arsenite and 150µg/ml antimycin-A for 4 hours after several trials.

### Generation of stable cell line overexpressing Alkbh1

B35 cells were cultured in a 96 well plate at density of 2 × 10^4^/well a day prior to the transfection. The next day GFP tagged Lenti-viral particles encoding Alkbh1 or Mock particles were seeded into each well at 5 × 10^4^, 12.5 × 10^4^ or 25 × 10^4^ transfection unites (TU) per well with 20µg Polybrene per well in full growth medium without antibiotics. Three days after transfection the transfection efficiency was evaluated using fluorescent microscopy to visualize the number of cells expressing GFP (Supplementary figure 4). Transfection efficiency was > 90% for all the concentrations used. Stable cell lines were generated using single colony expansion by passaging the cells into 96 well plate at a density of 1 cell per well. Colonies originating from cells expressing GFP were propagated and GFP intensity observed regularly to evaluate the degree of transfection and insure stable expression. Following colony expansion, Alkbh1 overexpression was evaluated using western blotting and compared to the Mock transfected cell line (Figure 4A). The Alkbh1 over-expressing cell line is referred to as B35^Alk^, while the Mock transfected cell line is referred to as B35^Mock^. Cells were exposed to stress using As (1mM) or antimycin (200µg/ml) for 4 hours.

### RNA extraction and northern blotting

Following stress experiments, cells were lysed using Qiazol (Qiagen, Hilden, Germany; Cat# 79306) and RNA extracted using miRNeasy mini kit (Qiagen, Cat# 217004) as described previously^2, 25^ and as per manufacturer protocol. RNA concentration was analyzed using nanodrop-one (Thermo Fischer).

For SYBR gold staining; Equal amounts of total RNA were mixed with sample buffer (Novex 2x TBE-urea sample buffer, Invitrogen, Cat# LC6876) and loaded onto Novex TBE-Urea Gels 10% (Cat# EC68755; Thermo Fisher) after denaturation by 70°C for 3 minutes and separated by gel electrophoresis using 10% TBE-Urea gels (Invitrogen, Cat# EC68755BOX) at 180V for 60 min in 1x TBE electrophoresis buffer (Bio-Rad, Cat# 161-0733). Gels were then incubated with SYBR gold fluorescent nucleic acid dye (Cat# S11494; Thermo Fisher) in 0.5x tris-borate-EDTA (TBE) for 40 min at room temperature and the images were acquired using ChemiDoc (Bio-Rad Laboratories).

1-methyadenosine (m^1^A) immunonorthern blotting (INB) and DIG-labeled probe-based northern blotting were performed as we previously described^2, 25, 29^. In brief, following electrophoresis, RNA was transferred onto Hybond N+ positive charged nylon membranes (Cat# RPN303B; GE Healthcare, Little Chalfont, UK) using semi-dry transfer and cross-linked using ultraviolet light. Membranes were either probed using anti-m^1^A antibody as per our published protocol^29^ or hybridized with 3’DIG-labeled probes against the 5’half of different tRNAs to evaluate the cleavage as we described previously^2^. The probes used in this study were:

For cytosolic tRNA:

5’tRNA-Arg^CCT^: 5’-AGGCCAGTGCCTTATCCATTAGGCCACTGGGGC

5’tRNA-Leu^AAG^: 5’-AATCCAGCGCCTTAGACCGCTCGGCCATGCTACC

5’tRNA-Gln^CTG^: 5’-AGTCCAGAGTGCTAACCATTACACCATGGAACC

5’tRNA-Gly^GCC^: 5’-GCAGGCGAGAATTCTACCACTGAACCACCAATGC

5’tRNA-Ala^AGC^: 5’-CTAAGCACGCGCTCTACCACTGAGCTACACCCCC

5’tRNA-Met^CAT^: 5’GACTGACGCGCTACCTACTGCGCTAACGAGGC

5’tRNA-Lys^CTT^: 5’GTCTCATGCTCTACCGACTGAGCTAGCCGGGC

5’tRNA-Glu^CTC^: 5’GCGCCGAATCCTAACCACTAGACCACCAGGGA

For mitochondrial tRNA:

5’mit-tRNA-Asn^GTT^: 5’-AGCTAAATACCCTACTTACTGGCTTCAATCTA

5’mit-tRNA-Tyr^GTA^: 5’-AGTCTAATGCTTACTCAGCCACCCCACC

5’mt-tRNA-Met^CAT^: 5’CCCGATAGCTTAGTTAGCTGACCTTACT

5’mt-tRNA-Gly^TCC^: 5’GTCAGTTGTATTGTTTATACTAAGGGAGT

5’mt-tRNA-Ala^TGC^: 5’ATCAACTGCTTTAATTAAGCTAAATCCTC

Sequences for Cytosolic tRNAs were acquired from GtRNAdb (http://gtrnadb.ucsc.edu/) and for mitochondrial tRNAs from mitotRNAdb (http://mttrna.bioinf.uni-leipzig.de/mtDataOutput/).

Following acquisition of images using the DIG-labeled probes, membranes were stripped using 0.1x SSC – 0.5% sodium dodecylsulphate (SDS) solution at 80°C for 30 minutes then washed briefly in PBS-T and re-probed or stored in 5x SSC buffer for long term storage at 4°C.

### Annexin V flowcytometry (FACS) assay

Apoptosis and cell death were analyzed using flowcytometry Annexin V assay as per our published protocol^2^. In brief, following exposure to stress in 24 well plate, cells were collected by trypsinization using 0.25% Trypsin in EDTA and washed using Annexin V binding buffer (0.5% bovine serum albumin (BSA) + 0.2mM Calcium Chloride in Fluorobrite DMEM medium [Gibco, Cat# A18967-01]). Then, cells were incubated with Alexa Fluor 647 Annexin V antibody (Biolegend, Cat# 640943) and Propidium Idodie (Invitrogen, Cat# P3566) for 15 minutes. The samples were analyzed using FACS canto II. Results were presented as fold-change (ratio) to unstressed controls.

### Cell growth analysis

For analysis of cell growth of Lenti-transfected cells, cells were cultured at a density of 1 × 10^6^ per well in a 6 well dish and the cell number was counted daily under microscopy using trypan blue staining (Takara, Japan. Cat# Y50015). Cells were also seeded at a low density in 96 well plate (5000 cells per well) and cultured for 7 days then cell growth was evaluated using WST-8 cell assay (Cell counting reagent SF, Nacalai tesque, Cat# 07553-44) under different growth conditions (Full growth medium, Serum free, glucose free or serum/glucose free growth media).

### Western blotting

Western blotting was performed as previously described^32^. Briefly, cells were collected and lysed in NE-PER nuclear and cytoplasmic extraction reagent (Thermo Fischer, Cat# 78833). The protein content was determined using the Brachidonic-acid assay kit (Thermo Fisher Scientific, Cat# 23227). Equal loads of proteins were separated with the Mini-PROTEAN TGX system (Bio-Rad Laboratories) and transferred to polyvinylidene difluoride membrane (Bio-Rad Laboratories). Membranes were incubated with antibody against Alkbh1 (1:500, Rabbit monoclonal, abcam, Cat# ab195376) overnight at 4□. After incubation with horseradish peroxidase-conjugated secondary antibody (IgG detector, Takara Clontech, Cat# T7122A-1), the antigen was detected using chemiluminescence Western blotting detection reagents (Pierce ECL Western Blotting Substrate, Thermo Fisher Scientific Inc., Rockford, IL, the USA, Cat# 32106). The image was scanned with ChemiDoc (Bio-Rad Laboratories). Membranes were striped using stripping buffer (Restore Western Blotting Stripping Buffer, Thermo Fisher Scientific Inc., Cat# 21059) and re-probed using anti-beta actin (1:5000, Rabbit IgG, Cell Signaling Technology, Cat# 13E5) using the same steps that were used previously. Semi-quantification was performed utilizing the ImageJ software after correcting the signal to an internal control (beta-actin).

### Statistical analysis

Statistical analysis was performed using SPSS v20 (IBM). All data are represented as mean ± standard deviation (SD). ANNOVA with Turkey post Hoc test was performed to compare between groups following FACS experiments. All experiments were repeated twice or thrice, and each experiment was performed with duplicate or triplicate samples for replicability assessment. No data was excluded from the analysis.

## Acknowledgment

The authors report no conflict of interest nor any ethical adherences regarding this work

## Author contribution

**Rashad:** Study conception and design, administration, developed protocols, conducted experiments, analyzed the data and wrote the manuscript. **Han:** Conducted experiments. **Sato:** conducted OGR-R experiments. **Niizuma:** Administration, Funding acquisition. **Mishima, Abe and Tominaga:** Critically revised the manuscript. **All authors:** approved the final version of the manuscript.

