## Supplementary Figures for "The stress specific impact of ALKBH1 on tRNA cleavage and tiRNA generation"

2-1 Seriyo-machi, Aoba-ku, Sendai, Japan.

Postal code: 980-8575

### Supplementary Figures

**Supplementary Figure 1:** Cropped gels in figure 1A

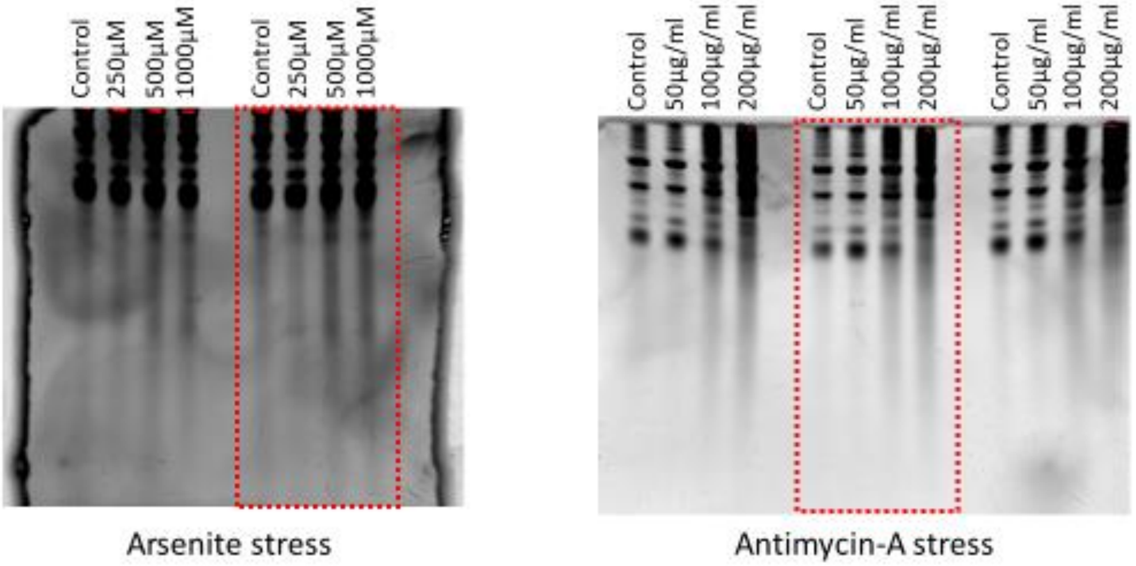

**Supplementary Figure 2:** Cropped gels in Figure 1B

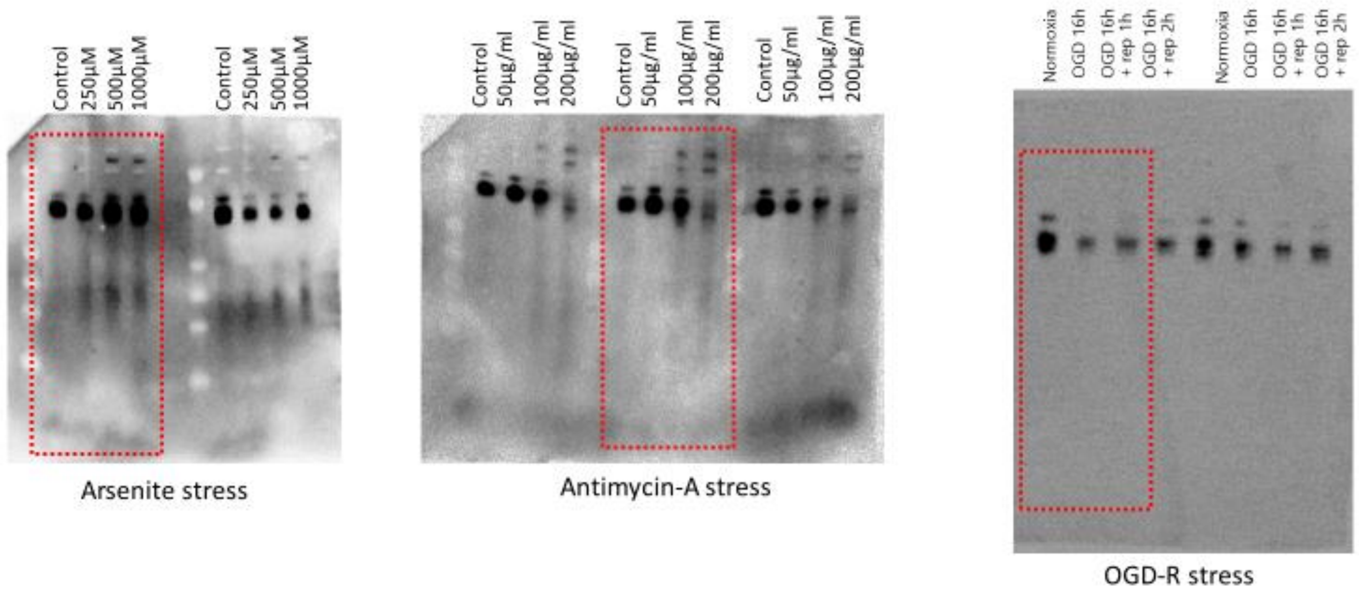

**Supplementary Figure 3:** Analysis of cell growth after Alkbh1 stable overexpression. **A:** WST-8 assay 1 week after seeding 5000 cells in 96 well plate and growth using different media (Medium 1: Full growth medium, Medium 2: Serum free medium, Medium 3: glucose free medium, Medium 4: serum/glucose free medium). **B:** Daily cell counting after platting $1x10^6$ in 6 well plate showing equal growth in $B35^{Mock}$ and $B35^{ALK}$ cells.

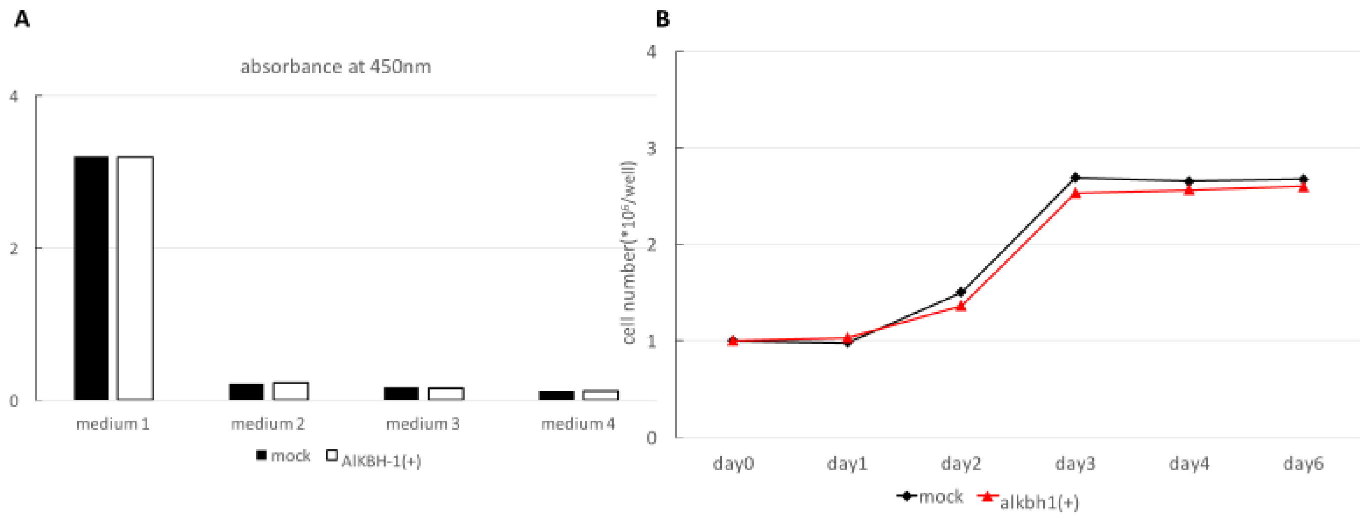

**Supplementary Figure 4:** GFP signal analysis 48 hours after transfection of B35 cells with Lentiviral particles encoding Alkbh1 or Mock sequence showing > 90% transfection efficiency

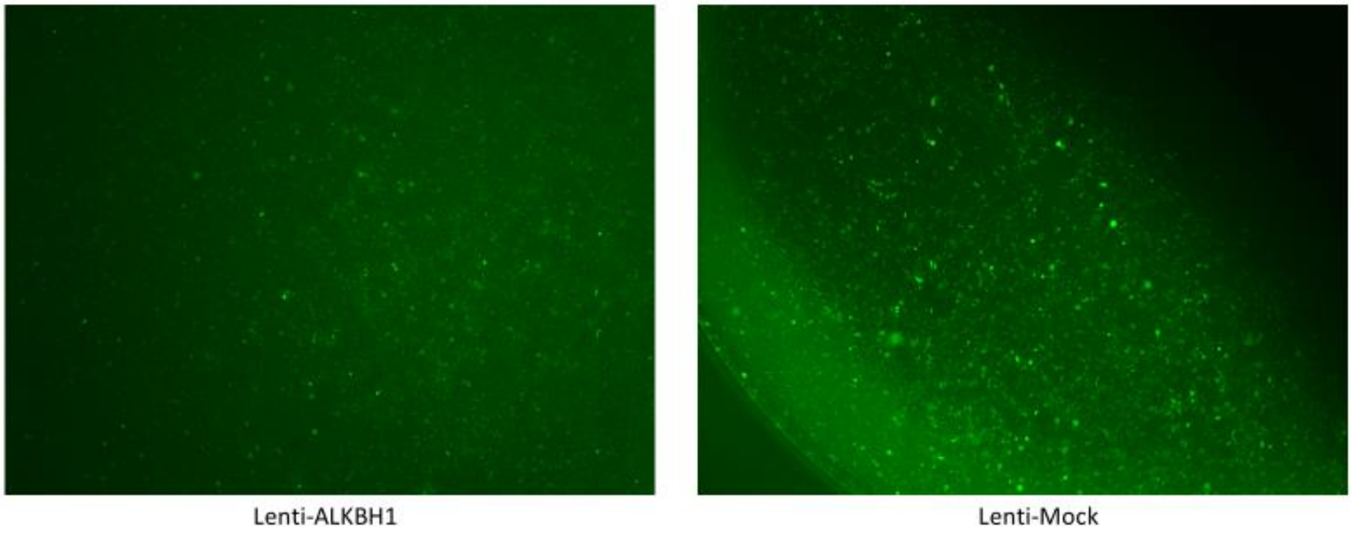
